# lionessR: single-sample network reconstruction in R

**DOI:** 10.1101/582098

**Authors:** Marieke L. Kuijjer, John Quackenbush, Kimberly Glass

## Abstract

We recently developed LIONESS (Linear Interpolation to Obtain Network Estimates for Single Samples), a method that can be used together with network reconstruction algorithms to extract networks for individual samples in a population. LIONESS was originally made available as a function within the PANDA (Passing Attributes between Networks for Data Assimilation) regulatory network reconstruction framework. In this application note, we describe lionessR, an R implementation of LIONESS that can be applied to any network reconstruction method in R that outputs a complete, weighted adjacency matrix. As an example, we use lionessR to model single-sample co-expression networks on a bone cancer dataset, and show how lionessR can be used to identify differential co-expression between two groups of patients.

**Availability and implementation:** The lionessR open source R package, which includes a vignette of the application, is freely available at https://github.com/mararie/lionessR.

## I. INTRODUCTION

Modeling and analyzing biological networks has become an invaluable tool in the analysis of genomic data. While gene expression profiles give us a snapshot of the state of a cell or tissue, network inference algorithms give an estimate of the extent to which genes or gene products interact [1]. Many network inference methods exist [2]. However, these are limited by the fact that they require population-level data but only infer one “aggregate,” or condition-specific, network.

We developed LIONESS, or Linear Interpolation to Obtain Network Estimates for Single-Samples [3], as a way of using population-level networks to estimate the corresponding network in each individual sample. LIONESS is based on the idea that each sample has its own network and that each edge in an aggregate network is the “average” (a linear combination) of that edge’s weight across these individual sample networks. LIONESS starts by modeling an aggregate network on an entire population and then removes one sample and rebuilds the network. LIONESS then compares the network with and without an individual sample, and uses a linear equation to estimate the network for the withheld sample:

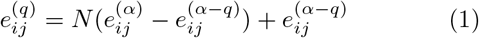

where 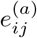 is the weight of an edge between nodes *i* and *j* in a network modeled on all (*N*) samples and 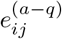 is the weight of that edge in a network modeled on all samples except the sample of interest (*q*). Thus, by sequentially leaving out each sample in a population, one can use LIONESS to estimate a network specific to each sample.

LIONESS network estimation is included as an option in our Python regulatory network reconstruction tool Py-Panda [4]. Here we present lionessR, a user-friendly R package to model single-sample networks for any network inference method.

## II. APPROACH

### II.1. The lionessR package

Within the lionessR package, the `lioness()` function applies LIONESS (Equation 1) to the output of a network reconstruction algorithm, as defined by the function `netFun`. The default network reconstruction algorithm in `netFun` is Pearson correlation, which builds co-expression networks by returning an adjacency matrix of Pearson correlation coefficients between pairs of genes. However, `netFun` can be substituted with any other uni-or bipartite network reconstruction algorithm that returns a complete, weighted adjacency matrix. `lioness()` returns an R data frame that includes weights for all edges in each of the sample-specific networks.

### II.2 Application of lionessR to a bone cancer dataset

As an example, we have performed an analysis applying `lioness()` to a gene expression dataset from 54 high-grade osteosarcoma biopsies [5] (Gene Expression Omnibus accession number GSE42352), which is included with the package. High-grade osteosarcoma is an aggressive primary bone tumor that has peak incidence in adolescents and young adults. About 45% of patients develop metastases, and most metastatic patients eventually die from the disease [6]. We performed a differential co-expression network analysis comparing short-versus long-term metastasis-free survival (MFS) to understand co-regulation differences between the groups and to search for potential therapeutic targets.

For this demonstration, we separated patients into two groups based on those who developed metastases within five years (n=19) and those who did not (n=35). These were the same groups analyzed by Buddingh et al. [7] to compare gene expression levels between short- and long-term MFS. For simplicity, we limited our analysis to the 500 most variable genes between groups. We used `lioness()` to model 54 single-sample networks based on Pearson correlation, one for each individual in the population, using the entire population to estimate the background network, with the code:

~~~
cormat <- lioness(dat, netFun),
~~~

where `dat` is the input expression data and `cormat` the lioness output.

We next asked whether there were differences in network edge weights between the short- and long-term MFS groups. To reduce the number of statistical tests on these networks, we modeled condition-specific networks and selected those edges that had an edge-weight difference of at least 0.5. We then performed a LIMMA analysis [8] to identify those edges whose weights differed significantly between the groups. In parallel, we also used LIMMA to test for significant differences in gene expression levels between groups. We visualized the 50 most significant edges (all nominal *p* < 0.001, *FDR* < 0.15) in a network diagram (Figure 1).

**Figure 1.**
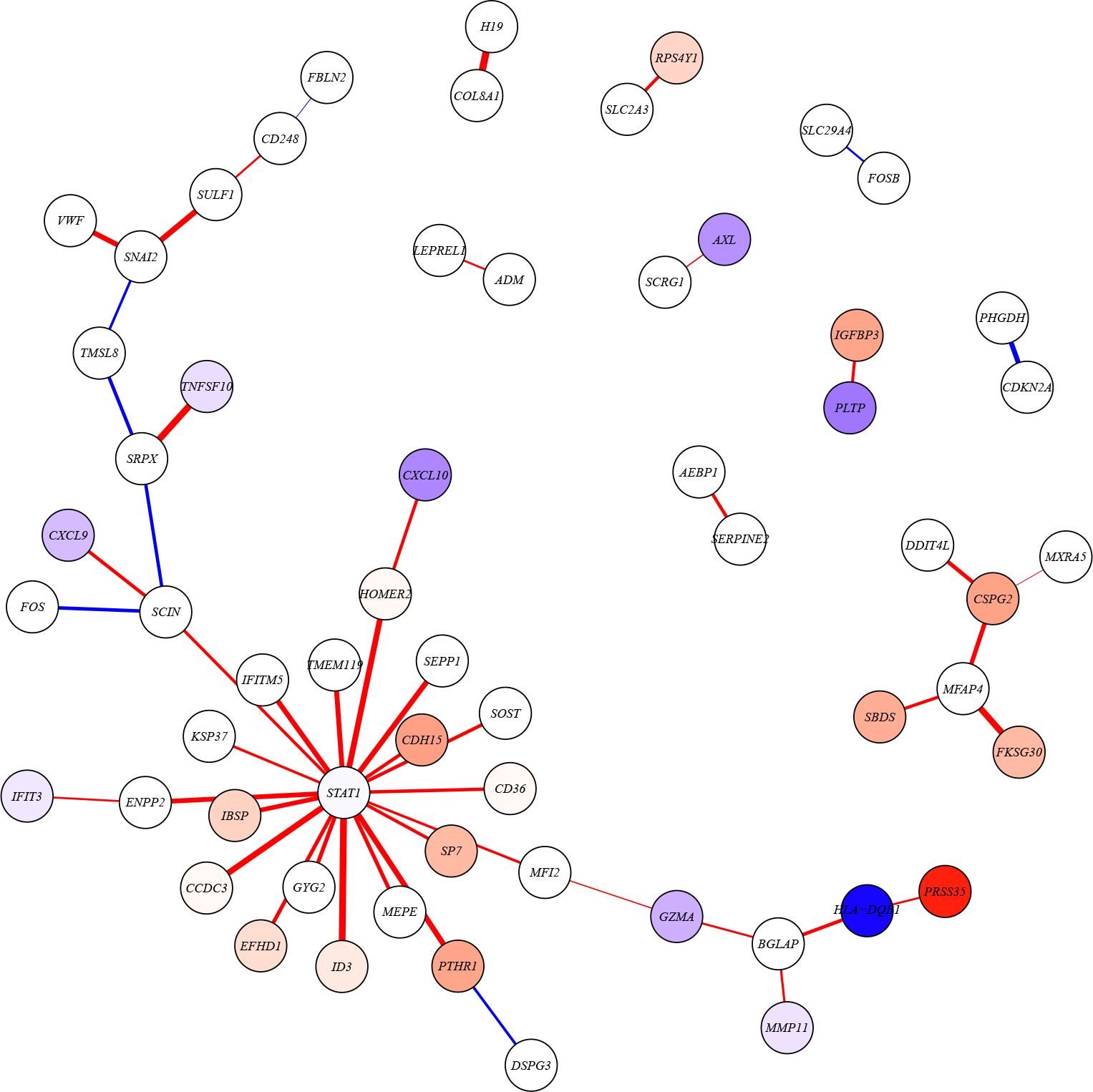
Network of the 50 most significant edges from a LIMMA analysis on single-sample co-expression edge weights between patients with poor and better MFS. Edges are colored based on whether they have higher weights in patients with poor (red) or better (blue) MFS. Thicker edges represent higher log fold changes. Nodes (genes) are colored based on the t-statistic from a differential expression analysis. Nodes with absolute t-statistic < 1.5 are shown in white, nodes in red/blue have higher expression in patients with poor/better MFS, respectively.

We identified a highly connected gene, or network “hub,” among the nodes connected to the top 50 edges— *STAT1*, or Signal Transducer And Activator Of Transcription 1. *STAT1* is a transcription factor and thus potentially differentially regulates the target genes with which it is correlated. In fact, all of the edges connected to *STAT1* had a moderate to strong negative correlation (range *R* = [−0.42, −0.84], median *R* = −0.67) in the samples with better MFS, whereas these edges had a weak to moderate positive correlation (range *R* = [0.14, 0.42], median *R* = 0.30) in the poor MFS group. This indicates that *STAT1* likely represses expression of these genes in patients with long-term MFS. However, this repression is lost in patients with short-term MFS. It has been previously shown that in tumors with good prognosis, high *STAT1* expression inhibits bone formation [9]. Many of the target genes we identified that connect to *STAT1* (Figure 1) are annotated to the Gene Ontology term “ossification,” consistent with this result.

More importantly, *STAT1* is a transcription factor in the interferon signaling pathway—a pathway known to be involved in osteosarcoma, and for which targeted treatment options are available [10]. This indicates that individual patient co-expression network analysis with lionessR can pinpoint potential candidates for personalized medicine.

## III. CONCLUSION

Network reconstruction algorithms can be used to better understand the interplay between different molecules in the cell. Until recently, it was not possible to model networks for individual samples or patients, which limited the usability of network methods, as aggregate networks do not allow modeling of heterogeneity in a population. We recently developed a method that can estimate individual sample networks by using linear interpolation to iteratively estimate networks for each individual in a population [3]. The lionessR package allows users to apply this method in combination with different network reconstruction algorithms, including Pearson correlation. As an example, we modeled single-sample networks based on the 500 genes with the highest variability in expression in an osteosarcoma dataset. We divided this dataset into two groups—patients with either short-term or long-term MFS. Comparing these two collections of networks using a LIMMA analysis, we identified *STAT1* to be significantly co-expressed with a set of “target” genes in biopsies of patients with poor survival. This set of genes was highly associated with biological processes important in osteosarcoma. In addition, *STAT1* is part of a biological pathway for which targeted treatment is available. This example highlights how single-sample co-expression network analysis can be used to inform us on potential precision medicine applications. The lionessR package is a user-friendly tool to perform such analyses.

